# Taxonomical and functional diversity of *Saprolegniales* in Anzali lagoon, Iran

**DOI:** 10.1101/860429

**Authors:** Hossein Masigol, Seyed Akbar Khodaparast, Reza Mostowfizadeh-Ghalamfarsa, Keilor Rojas-Jimenez, Jason Nicholas Woodhouse, Darshan Neubauer, Hans-Peter Grossart

## Abstract

Studies on the diversity, distribution and ecological role of *Saprolegniales* (*Oomycota*) in freshwater ecosystems are currently receiving attention due to a greater understanding of their role in carbon cycling in various aquatic ecosystems. In this study, we characterized several *Saprolegniales* species isolated from Anzali lagoon, Gilan province, Iran, using morphological and molecular methods. Four species of *Saprolegnia* were identified, including *S. anisospora* and *S. diclina* as first reports for Iran. Evaluation of the ligno-, cellulo- and chitinolytic activities were also measured using plate assay methods. Most of the *Saprolegniales* isolates were obtained in autumn and nearly 50% of the strains showed chitinolytic and cellulolytic activities. However, only a few *Saprolegniales* strains showed lignolytic activities. This study has important implications for better understanding the ecological niche of oomycetes, and to differentiate them from morphologically similar but functional different aquatic fungi in freshwater ecosystems.

## Introduction

*Saprolegniales*, as a monophyletic order, belong to the phylum *Oomycota*. It includes biflagellate heterotrophic microorganisms that have eucarpic mycelial and coenocytic thalli of unlimited growth. They produce asexual (sporangia) and sexual (gametangia) structures delimited by septa. *Saprolegniales* are mainly predominantly freshwater saprophytes of plant and animal debris (Beakes and Sekimoto 2009). This order contains three families; *Achlyaceae* (four genera), *Saprolegniaceae* (11 genera) and *Verrucalvaceae* (7 genera) (Beakes et al. 2014; Molloy et al. 2014; Beakes and Thines 2017; Rocha et al. 2018). Among these, *Achlya, Brevilegnia, Dictyuchus, Leptolegnia, Plectospira, Saprolegnia* and *Thraustotheca* have been commonly reported to inhabit freshwater ecosystems (Czeczuga et al. 2005; Mousavi et al. 2009; Marano et al. 2011).

*Saprolegniales* species have been recently receiving increased attention due to their wide distribution, ubiquitous occurrence (Liu and Volz 1976; Kiziewicz and Kurzątkowska 2004; Nascimento et al. 2011), their devastating fish pathogenicity in aquaculture and fish farms and responsibility for massive decline of natural salmonid populations (Griffiths et al. 2003; Van West 2006; Romansic et al. 2009; Van Den Berg et al. 2013). Aside from their pathogenicity, many authors have also investigated relative frequencies of *Saprolegniales* throughout different seasons in relation to physico-chemical features of the respective freshwater ecosystems (El-Hissy and Khallil 1991; Czeczuga et al. 2003; Paliwal et al. 2009). Although these oomycetes are generally isolated from plant debris, their involvement in organic matter degradation in freshwater ecosystems remains less clear.

Species identification of *Saprolegniales* is largely based on morphological features (Coker 1923; Seymour 1970; Johnson et al. 2002). However, this identification is perplexing due to several reasons. First of all, in many cases, morphological and morphometric characters are vague and variable. Secondly, more determinative features like sexual structures are not always produced *in vitro*. Also, lack of type species and accurate description make it even harder to define specific species (Sandoval-Sierra et al. 2015, 2014). Recently, sequencing of the ribosomal internal transcribed spacer (ITS) has been applied to create a phylogenetic framework within which to address issues of morphological and taxonomic ambiguity (Steciow et al. 2014). Whether complementary molecular targets, in addition to the *de facto* ITS region, would improve this approach or are even necessary is still open to debate (Robideau et al. 2011).

In this study, we investigated the diversity and seasonality of various strains of *Saprolegniales* isolated from Anzali lagoon, Iran. In total, we obtained 511 isolates from three locations during 2017 and studied their seasonality. From these, 23 isolates were randomly selected representing different sampling time points and locations and identified using morphological and phylogenetic analyses. In addition to their taxonomy, we tested the hypothesis raised by Masigol et al. (2019) that fungi and *Saprolegniales* differ in their affinity for polymeric dissolved organic matter (DOM) and consequently in their involvement in aquatic DOM degradation and cycling. To this, we evaluated the ligno-, cellulo- and chitinolytic activities of the selected strains. Our results have important implications for understanding the different roles of fungi and *Saprolegniales* in aquatic ecosystems.

## Materials & Methods

### Sampling site

Anzali lagoon is situated at the Caspian Sea near Bandar-e Anzali, in the northern Iranian province of Gilan. The lagoon divides Bandar-e Anzali into two parts, and is home to both the Selke Wildlife Refuge and the Siahkesheem Marsh. Three sampling sites as representatives of the main habitats in Anzali lagoon were selected:1) river entrance, 2) shallow water habitat and 3) urban habitat (Fig. 1).

**Fig.1.**
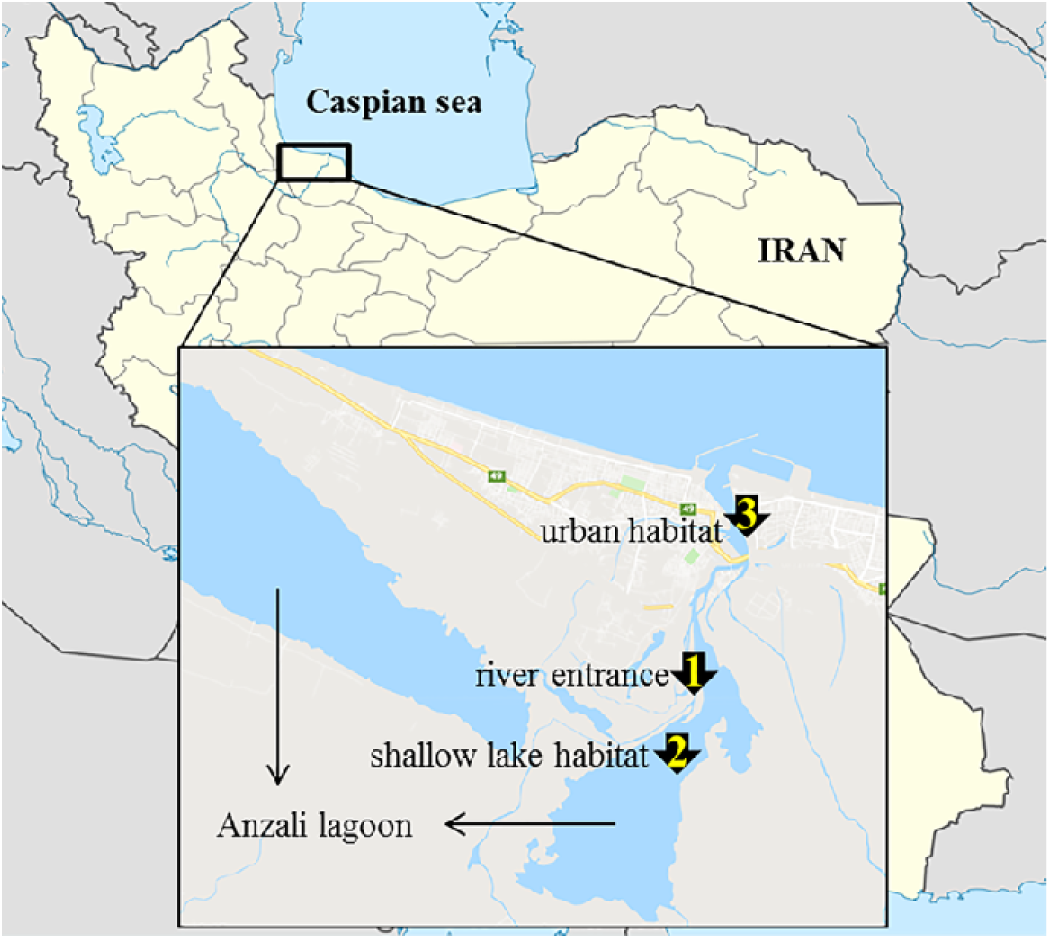
Sampling locations (1: river entrance, 2: shallow lake habitat and 3: urban habitat) for isolation of *Saprolegniales* taxa from Anzali lagoon, Iran during 2017

### Seasonal distribution of *Saprolegniales* and isolation

Throughout 2017, 511 *Saprolegniales* isolates were isolated using the methods described earlier by Coker (1923) and Seymour (1970). In brief, samples of decaying leaves of the dominant local vegetation collected from the three sampling locations were brought to the mycology laboratory of the University of Guilan in separate sterile polyethylene bags. Leaves were cut into approximately ten equal pieces (0.5×0.5 cm). After washing with distilled water, they were incubated at 20-25°C in sterilized plates containing 10 mL sterile distilled water with 20 sterilized hemp seed halves (*Cannabis sativa* L.) (Middleton 1943). Temperature and pH of surface water were continuously recorded immediately after collecting decaying leaves. Three replicates were considered for each location. The average number of colonized hemp seed halves from 10 Petri dishes was used to estimate the abundance of *Saprolegniales* throughout the year. The presence of *Saprolegniales* was confirmed by observing at least one of the general features of oomycetes such as oogonia, sporangia, large and aseptate mycelia and motile zoospores.

After three to five days, a piece of mycelia from the colonized hemp seed halves was transferred to a fresh CMA-PARP medium (CMA-PARP; 40 g/L ground corn meal, 0.5 g/L ampicillin, 0.01 rifampicin, 0.2 g/L Delvocidsalt and 0.1 g/L Pentachloronitrobenzene (PCNB), 15 g/L agar) (Kannwischer and Mitchell 1981). This step was repeated three to five times to achieve bacterial free (axenic) cultures. A single hypha was transferred to cornmeal agar (CMA; 40 g/L ground corn meal, 15 g/L agar) (Seymour and Fuller 1987). The hyphal-tip technique was conducted three to five times to obtain a pure culture in CMA. The specimens of these new strains were then deposited in the Fungal Herbarium of the Iranian Research Institute of Plant Protection, Iran.

### Characterization of morphological features

Asexual and sexual structures of isolates were characterized and measured in liquid (water) cultures (n=30). To investigate strains failing to produce any sexual structures, several treatments were used. The nutrition treatments included (1) reciprocal culturing of all strains with one another and *Trichoderma* sp. (Brasier et al. 1978) on CMA, (2) hemp seed agar (HSA; 60 g/L ground hemp seeds, 15 g/L agar) (Hendrix 1964), (3) soybean agar (SA; 100g/L ground soybean seeds, 15 g/L agar) (Savage et al. 1968), (4) rape seed extract agar (REA; 100g/L ground rape seeds, 15 g/L) (Satour 1967), (5) carrot juice agar (CJA; 250 g/L boiled carrot extract, 20 g/L agar) (Ershad 1971), (6) mPmTG (2, 0.4, 0.4 and 12 g/L glucose, tryptone, peptonized milk and agar, respectively) (Moreau and Moreau 1936b), (7) immersing colonized CMA in glycerin (4%) (Moreau and Moreau 1936a) and (8) culturing the isolates in Petri dishes containing 10 boiled hemp seeds in distilled lake water and distilled water (50/50). The temperature treatments also included culturing the isolates in 5, 10, 15, 20, and 25°C in Petri dishes containing ten boiled hemp seeds in distilled lake water and distilled water (50/50).

### DNA extraction and PCR

The DNA extraction was conducted based on a slightly modified protocol of Montero-Pau et al. (2008). Briefly, 100 µL of alkaline lysis buffer (25 mM NaOH, 0.2 mM disodium EDTA, pH 8.0) was aliquoted into 1.5 mL tubes. Malt extract broth (MEB; 17 g/L malt extract) (Galloway and Burgess 1937) was used for growth of isolates. Mycelial mass was then transferred to the tube and centrifuged for 30 minutes at 9000 rpm. The tube was incubated at 95°C for 30 minutes and then cooled on ice for five minutes. Finally, 100 µL of neutralizing solution (40 mM Tris-HCl, pH 5.0) was added to the tubes. The final solution was vortexed and kept at −20°C. Nuclear ITS region was amplified in a Flexible PCR Thermocycler (Analytikjena, Germany) using ITS1/ITS4 (White et al. 1990) primers. Thermocycler program for amplification of the ITS region was: 94°C for 2 min for initial denaturation followed by 32 cycles of 94°C for 15 s, 53°C for 15 s, 72°C for 30 s, and a final extension at 72°C for 5 min (White et al. 1990). The resulting sequences were quality controlled using the Bioedit software (Hall et al. 2011) and submitted to GenBank (National Center for Biotechnology Information; http://www.ncbi.nlm.nih.gov) database (for accession numbers see Table1).

**Table 1.**
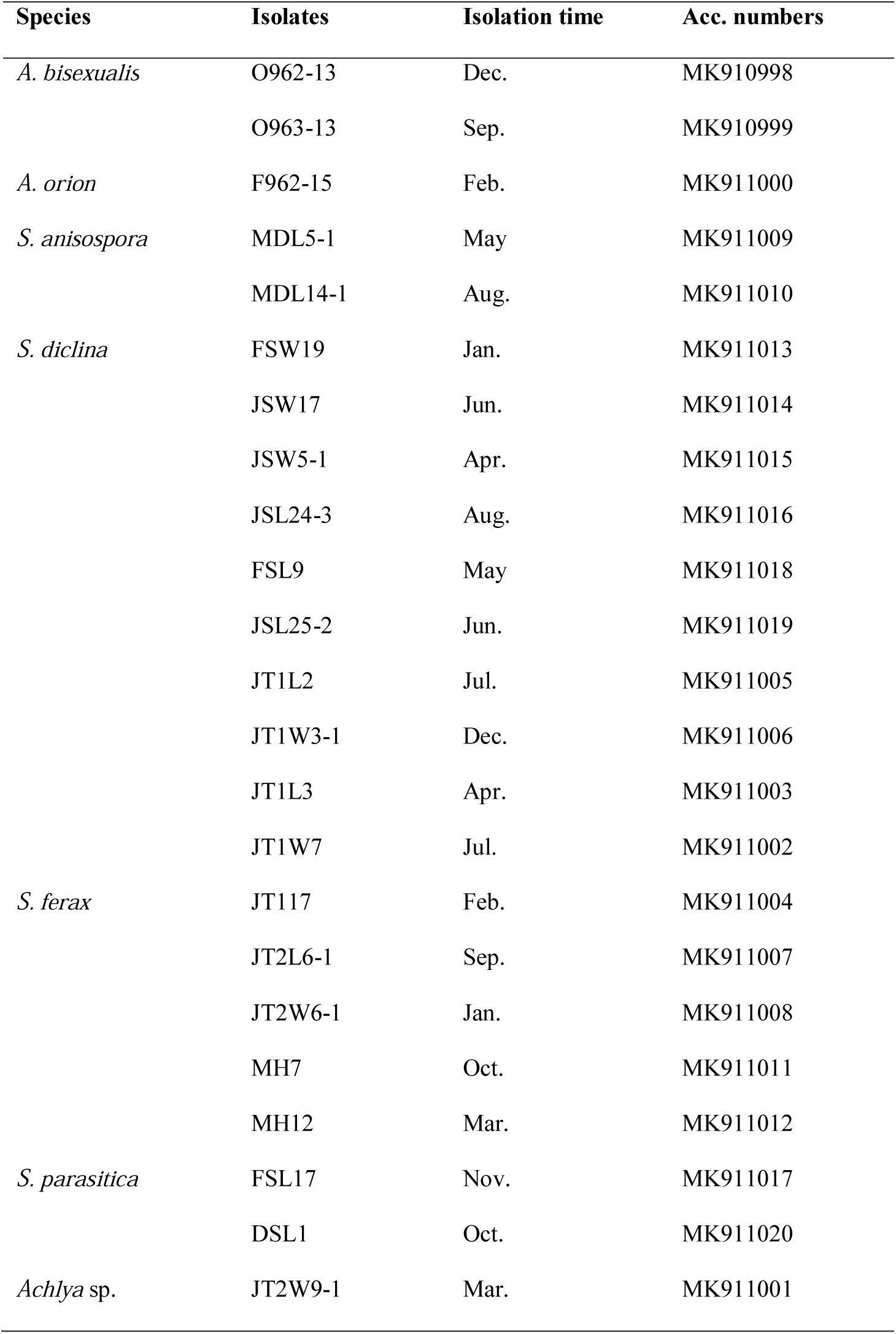
*Saprolegniales* isolates obtained from three coordinates Anzali lagoon (Rasht County, Iran) during 2017 and their GenBank accession numbers (ITS region).

### Phylogenetic analysis

Molecular taxonomical assignment of the isolates was performed by comparing sequences against the NCBI’s GenBank sequence database using the BLASTN search method. We selected and downloaded sequences of cultured isolates for performing phylogenetic analyses. The ITS sequences were aligned by the ClustalW algorithm (Thompson et al. 1994) and the phylogenetic tree was obtained using MEGA version 7 by the Maximum Likelihood (ML) method based on the Tamura-Nei model (Kumar et al. 2016).

### Screening for lignolytic, cellulolytic and chitinolytic activities

#### Lignolytic assay

Mycelia from the edge of 7-15 days cultures were transferred into 6-well plates containing the cultivation medium proposed by Rojas-Jimenez et al. (2017) (0.94 g KH_2_PO_4_, 1.9 g K_2_HPO_4_, 1.6 g KCl, 1.43 g NaCl, 0.15 g NH_4_Cl, 0.037 g MgSO_4_, 0.1 g yeast extract, 10 g malt extract, 15 g/L agar, pH 7.0) and mPmTG agar medium amended with one of the following substrates: (1) 0.1% wt/vol 2,20-Azino-bis 3-ethylbenzothiazoline-6-sulfonic acid diammonium salt (ABTS), (2) 0.02 and 0.005% wt/vol Bromocresol Green (BG), (3) 0.02 and 0.005% wt/vol Congo Red (CR), (4) 0.02 and 0.005% wt/vol Malachite Green (MG) (5) 0.02 and 0.005% wt/vol Phenol Red (PhR) (6) 0.02 and 0.005% wt/vol PolyR-478 (PR) (pH 5 + 7) (7) 0.02 and 0.005% wt/vol Remazol Brilliant Blue (RBBR), and (8) 0.02 and 0.005% wt/vol Toluidine Blue (TB) (Pointing 1999; Swamy and Ramsay 1999; Moreira et al. 2000; Novotny et al. 2001; Gill et al. 2002; Rojas-Jimenez et al. 2017). The concentration of different dyes in previous experiments is highly variable. Therefore, two different concentrations were used to ensure that the applied concentration does not impact on the growth or the enzymatic activities of the tested strains. The capacity of each strain to produce lignolytic activity was determined by decolorization of the aforementioned substrates in the area around the mycelia or as a colour change of the media after three weeks. We evaluated 1-33, 33-66, and 66-100% decolorization of the medium in the Petri dishes as weak, medium, and strong activity, respectively.

#### Cellulolytic assay

The same media as used for evaluation of lignolytic activities were amended with the following enzymatic carbon sources to investigate cellulolytic and pectolytic activities: (9) 7.5 g carboxymethylcellulose (CMC), (10) 7.5 g Avicel (AVL) and (11) 5g D-cellobiose (DCB) (Wood and Bhat 1988; Pointing 1999; Yoon et al. 2007; Jo et al. 2010). After three weeks of incubation, Congo Red (1 mg ml^-1^) was amended to the medium and incubated at room temperature for 15 min. Subsequently, the medium was rinsed with distilled water, and 30 mL of 1 M NaCl added. Degradation of CMC, Avicel, and D-cellobiose was confirmed by a transparent appearance of the medium (and mycelia) (Teather and Wood 1982; Pointing 1999).

#### Chitinolytic assay

The method proposed by Agrawal and Kotasthane (2012) was used to evaluate the chitinolytic properties of the isolated strains. Crab shell flakes were ground in a mortar and sieved through the top piece of a 130 mm two-piece polypropylene Buchner filter. Twenty grams of the sieved crab shell flakes were then treated with 150 mL of ∼12M concentrated HCl which was added gently and continuously stirred for 45 minutes under a chemical fume hood. The final mixture was passed through eight layers of cheese cloth to remove large chitin chunks. The product was treated with two litres of cold distilled water and incubated overnight under static conditions at 4°C. Sufficient amount of tap water was then passed through the product until the pH of the product reached 7.0. The final product was squeezed between coffee paper and then sterilized by autoclaving at standard temperature and pressure (STP) (15 psi, 20 minutes, 121°C) (Murthy and Bleakley 2012). The chitinase detection medium consisted of a basal medium comprising (per litre) 0.3 g of MgSO_4_.7H_2_O, 3.0 g of (NH_4_)_2_SO_4_, 2.0 g of KH_2_PO_4_, 1.0 g of citric acid monohydrate, 15 g of agar, 200 μL of Tween-80, 4.5 g of colloidal chitin (CC) and 0.15 g of Bromocresol Purple; the pH was adjusted to 4.7, and the neutralized medium autoclaved.

For each isolate, the experimental procedures mentioned above were repeated twice with three replicates each. When a positive result was observed, this was confirmed in a third experiment using tissue culture plates (TPP^®^) with the medium described above. We observed no conflicting results from any of the three experimental instances, for each of the tested isolates and substrates. Strains already tested by Rojas-Jimenez et al. (2017) were considered as positive controls.

#### Statistical analyses

We assessed whether there was a significant impact of month or broadly season at each of the three locations on the frequency of *Saprolegniales* colonisation of hemp leaves, using a two-way ANOVA. The relative contribution of temperature and pH to any spatio-temporal trends were assessed using Spearman’s Rank correlations. A propensity of individual genera or taxa to metabolise invidual substrates was assessed using a two-way ANOVA. All statistical analyses were conducted using GraphPad Prism version 8.1. (GraphPad Software, CA, USA).

## Results

### Relative abundance of *Saprolegniales* isolates

From all the oomycetes isolated from the three locations sampled along the year, 511 out of 720 (∼71%) were assigned to the order *Saprolegniales.* The relative abundance of *Saprolegniales* was higher at cold temperatures (autumn, winter and spring seasons) than in summer. The highest temperatures were recorded in Jul. 23th-Aug. 22th (in average∼33°C) and the lowest in Jun. 21th-Feb. 19th (in average∼3°C). The number of isolates from river entrance, shallow lake and urban habitats was negatively correlated with temperature (R^2^= 0.7233, 0.5047 and 0.7623, respectively). However, pH was constant and no correlation was observed for river entrance, shallow lake and urban habitats (R^2^ = 2E-05, 0.0037 and 0.0684, respectively). Of the 720 hemp seeds, only 209 were not colonized by *Saprolegniales* isolates (∼29%). These were either colonized by other microorganisms such as fungi and protists, or remained intact. Co-colonization of *Saprolegniales* isolates and other unwanted subjects or organisms was not counted as a positive result (Table 2).

**Table 2.**
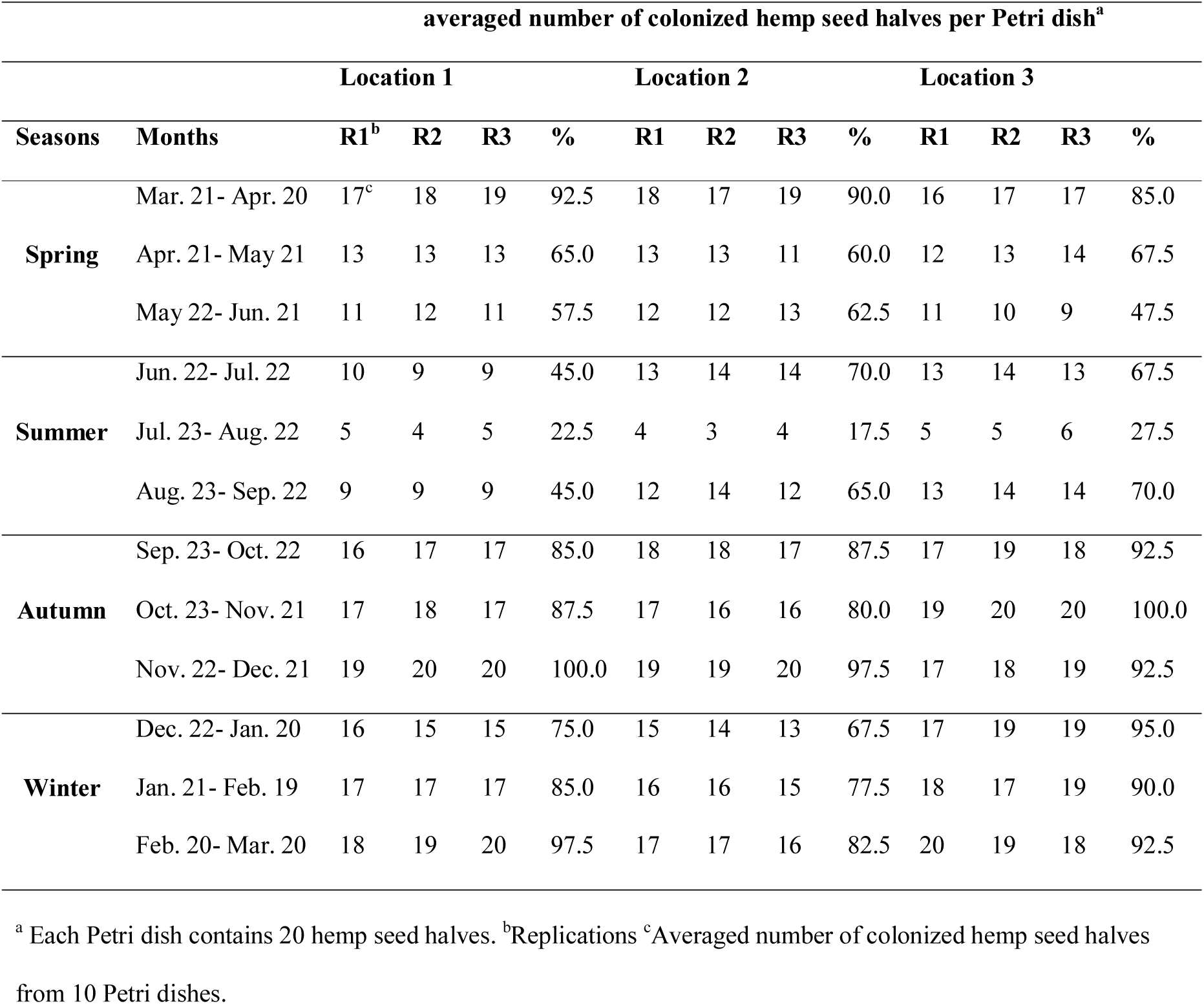
Number and percentage of colonized hemp seed halves per Petri dish isolated from three sampling sites in Anzali lagoon, Iran throughout 2017.

### Morphological and molecular identification

Of the selected isolates, 19 belonged to the genus *Saprolegnia* and four to the genus *Achlya*. Four isolates of *Achlya* failed to produce sexual structures under any circumstances and thus were considered as *Achlya* spp. (Isolates JT2W9-1, F962-15, O962-13 and O963-13). Morphology-based taxonomy was confirmed by phylogenetic analysis of ITS sequences of nrDNA inferred from maximum likelihood method.

#### Saprolegnia anisospora

(Pringsheim) de BaryBot. Zeitung (Berlin) 41:56. 1883 (Fig. 2a-c) Mycelium dense; main hyphae branched, hyaline to dark, with 16–46 μm (average 26 µm) width. Sporangia very abundant, mainly fusiform, straight, sometimes curved, renewed in cymose fashion, 80–405 ×18–50 μm (average 220×26 µm). They discharged spores and behaved as saprolegnoids. Cysts 9–13 μm in diameter (average 10 µm). Gemmae absent. Oogonia terminal, always spherical, always immature, 78–107 μm in diameter. Oogonial wall smooth. Oogonial stalks 1-3 times the diameter of oogonium, slender, slightly irregular and unbranched. Oospores never produced. No specific pattern was observed for any of our strains on CMA.

**Fig. 2.**
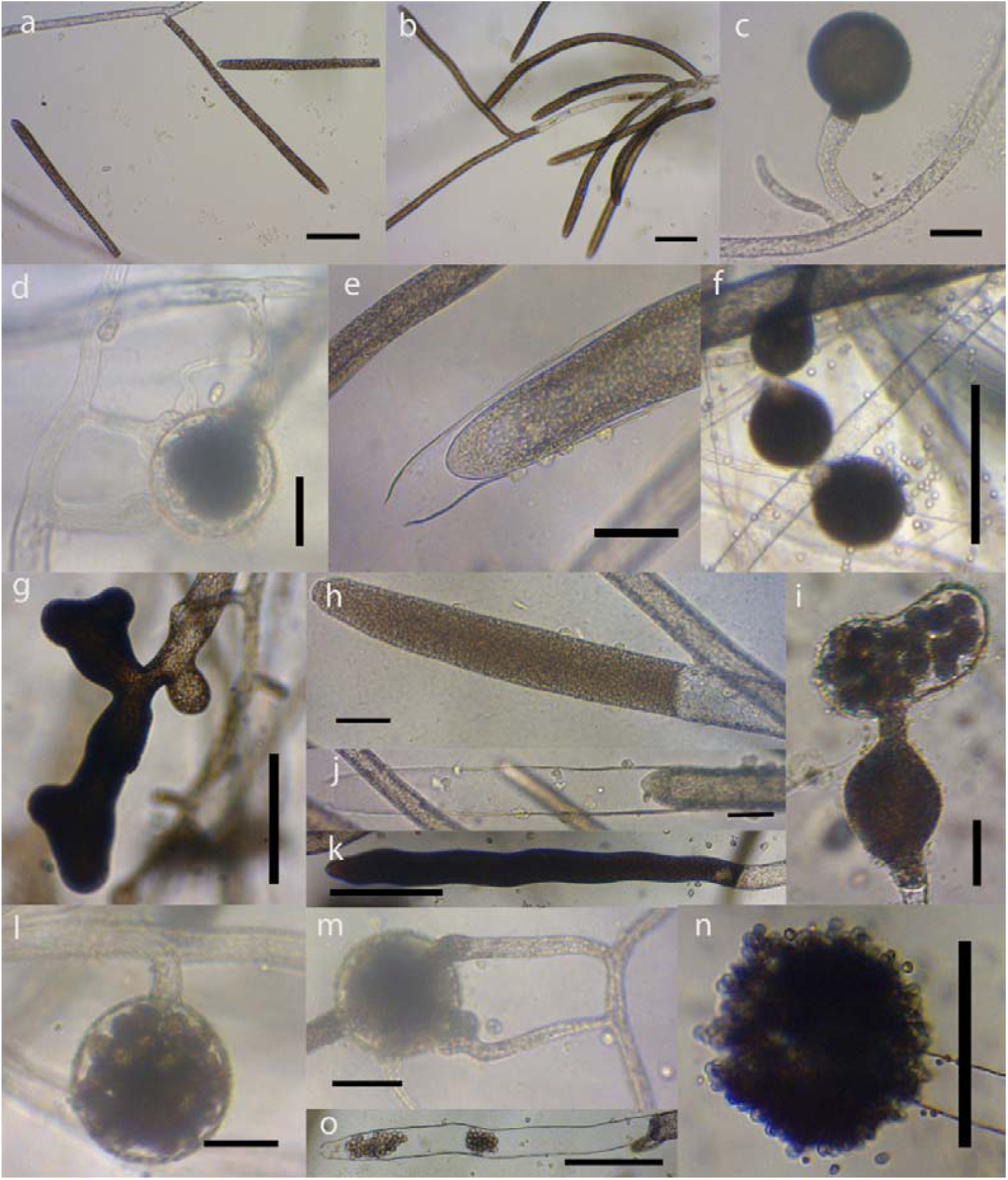
General morphological characteristics of selected isolates used in this study observed in water culture at room temperature; (a-c) fusiform straight and sporangia (a), cymose fashion renewal of sporangia (b) and terminal immature oogonia with stalk (c) of *Saprolegnia anisospora;* (d-f) diclinous and androgynous antheridia (d), internal renewal of sporangia and catenulate spherical gemma(f) of *S. diclina*; (g-i) an extremely irregular gemma (g), cylindrical sporangia (h) and an oogonium with irregular shape (i) of *S. ferax*; (j and l-m) internal renewal of sporangia (j), lateral oogonia and short stalk (l), diclinous antheridia (m) of *S. parasitica*, (k) sporangia, (o) empty sporangia and (n) spore clump of *Achlya*spp. (bar=50 µm, except for k, n and o which are 200 µm)

### Material examined

strains MDL5-1 and MDL14-1, on rotten leaves, Anzali lagoon, Anzali, Guilan, Iran, 10-8-2017, H. Masigol; GenBank Acc. No: ITS – MK911009 and MK911010.

#### Saprolegnia diclina

Humphrey Trans. Amer. Phil. Soc. (N.S.) 17:109, pl. 17 (Fig. 2d-f) Mycelium sparingly to moderately branched, 17–42 μm (average 35 µm) in width. Sporangia abundant, cylindrical, always straight, renewed internally, 120–976 ×18–69 μm (average 460×52 µm). They discharged spores and behaved as saprolegnoids. Cysts 8–11 μm in diameter (average 9 µm). Gemmae spherical, 66–110 μm in diameter, terminal, sometimes catenulate. Oogonia terminal, spherical, obpyriform, 75–105 μm in diameter. Oogonial wall smooth. Oogonial stalk 1–3 times the diameter of the oogonium, slender and unbranched. Oospores centric, spherical, 6–26 per oogonium and 14–28 μm in diameter. Antheridia abundant, diclinous and androgynous. No specific pattern observed for any of our isolates on CMA.

### Material examined

isolates JSL25-2, FSL9, JSL24-3, JSW5-1, JSW17 and FSW19, on rotten leaves, Anzali lagoon, Anzali, Guilan, Iran, 10-8-2017, H. Masigol; GenBank Acc. No: ITS – MK911019, MK911018, MK911016, MK911015, MK911014 and MK911013.

#### Saprolegnia ferax

(Grith.) Thuret Ann. Sci. Nat. Bot. 14:229 et spp. pl. 22. 1850 (Fig. 2g-i) Mycelium dense; main hyphae highly branched, hyaline to dark, 15–62 μm (average 44 µm) in width. Sporangia abundant, cylindrical, rarely fusiform, always straight, renewed sympodially, 60–490 ×22– 78 μm (average 352×51 µm). They discharged spores and behaved as saprolegnoids. Cysts 7–11 μm in diameter (average 10 µm). Gemmae were overabundant, extremely irregular, terminal and intercalary. Oogonia terminal, sometimes intercalary, spherical, obpyriform, spherical, 76–99 μm in diameter and sometimes with irregular shapes. Oogonial smooth. Oogonial stalk 1–3 times the oogonium diameter, slender and unbranched. Oospores centric, spherical, 2–48 per oogonium, 15–63 μm in diameter. Antheridia very rare, when present diclinous, 1–5 per oogonia. No specific pattern observed for any of isolates on CMA.

### Material examined

isolates JT1L3, JT1W7, JT117, JT1L2, JT1W3-1, JTL6-1 and JT2W6-1, on rotten leaves, Anzali lagoon, Anzali, Guilan, Iran, 10-8-2017, H. Masigol; GenBank Acc. No: ITS – MK911003, MK911002, MK911004, MK911005, MK911006, MK911007 and MK911008.

#### Saprolegnia parasitica

Coker *emend.* Kanouse, Mycologia 24:447, pls. 13. 1932 (Fig. 2j and l-m) Mycelium moderately branched, with 18–56 μm (average 39 µm) width. Sporangia cylindrical, renewed internally, 150–460 ×20–66 μm (average 325×52 µm). They discharged spores and behaved as saprolegnoid. Cysts 10–12 μm in diameter (average 11 µm). Gemmae absent in water cultures, when present they were terminal, spherical and single. Oogonia mainly lateral, sometimes terminal, 86–110 μm in diameter, very rarely catenulate. Oogonial wall smooth. Oogonial stalk very short, 0.4-0.9 times the oogonium diameter, slender, unbranched and sometimes absent. Oospores centric, spherical, 8–32 per oogonium and 12–29 μm in diameter. Antheridia diclinous. No specific pattern observed for any of our isolates on CMA.

### Material examined

isolates DSL1 and FSL17, on rotten leaves, Anzali lagoon, Anzali, Guilan, Iran, 10-8-2017, H. Masigol; GenBank Acc. No: ITS – MK911020 and MK911017.

### Phylogenetic analyses

The phylogenetic classification of ITS sequences (694 bp) of the different genera belonging to *Saprolegniaceae* (*Saprolegniales, Oomycota*), inferred from both maximum likelihood method was consistent with the morphology-based taxonomy. Moreover, we found for most morphological identified species some reliable sequences with high similarity (99-100%), so that these sequences clustered with our identified species in the same clade (Fig. 3).

**Fig. 3.**
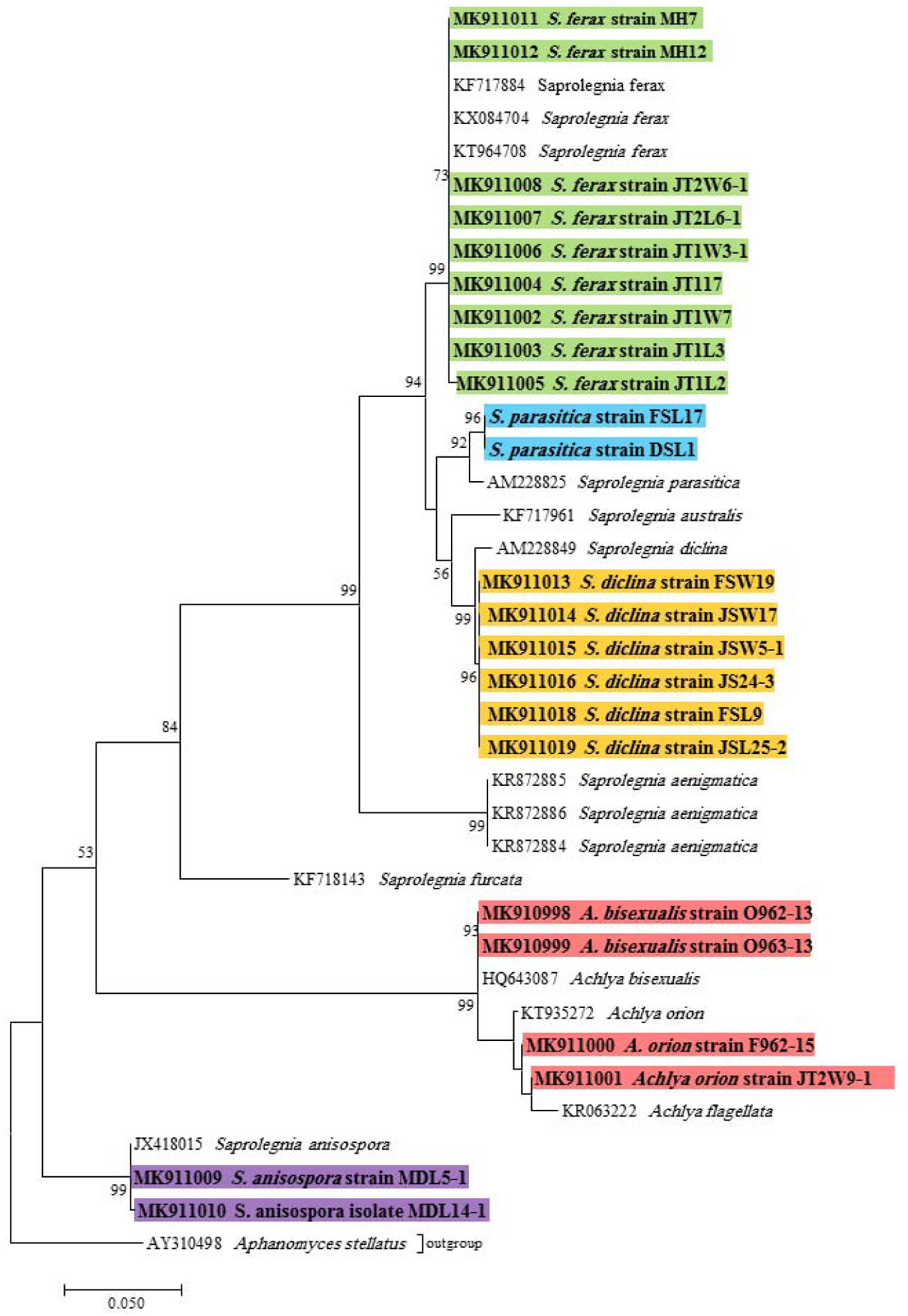
Phylogenetic tree of the family *Saprolegniaceae* (*Saprolegniales, Oomycota*). The analysis was performed based on alignment of the ITS1-5.8S-ITS2 region (694bp) using the maximum likelihood method from isolates in this study (bold and colored) and valid sequences from GenBank. Numbers next to the branches shows bootstraps values ≥ 50%.*Aphanomyces stellatus* was considered as outgroup

### Screening for lignolytic, cellulolytic, pectolytic and chitinolytic activities

Of all tested isolates, 61% showed chitinolytic activities. In addition, 52, 43, and 48% of all tested isolates showed cellulolytic activities in the medium amended with avicel (AVL), carboxymethylcellulose (CMC) and D-cellobiose (DCB), respectively. In contrast, no significant lignolytic activities were observed. Decolorization of dyes was detected only in the medium amended with Bromocresol Green (BG) and Toluidine Blue (TB) (39 and 8% of isolates, respectively). In some cases, isolates failed to grow, especially in the medium amended with Phenol Red (phR). Also, in media amended with Congo Red, adsorption by mycelia was observed. Thus, it was not considered as a proof for lignolytic activity (table 3).

**Table 3.**
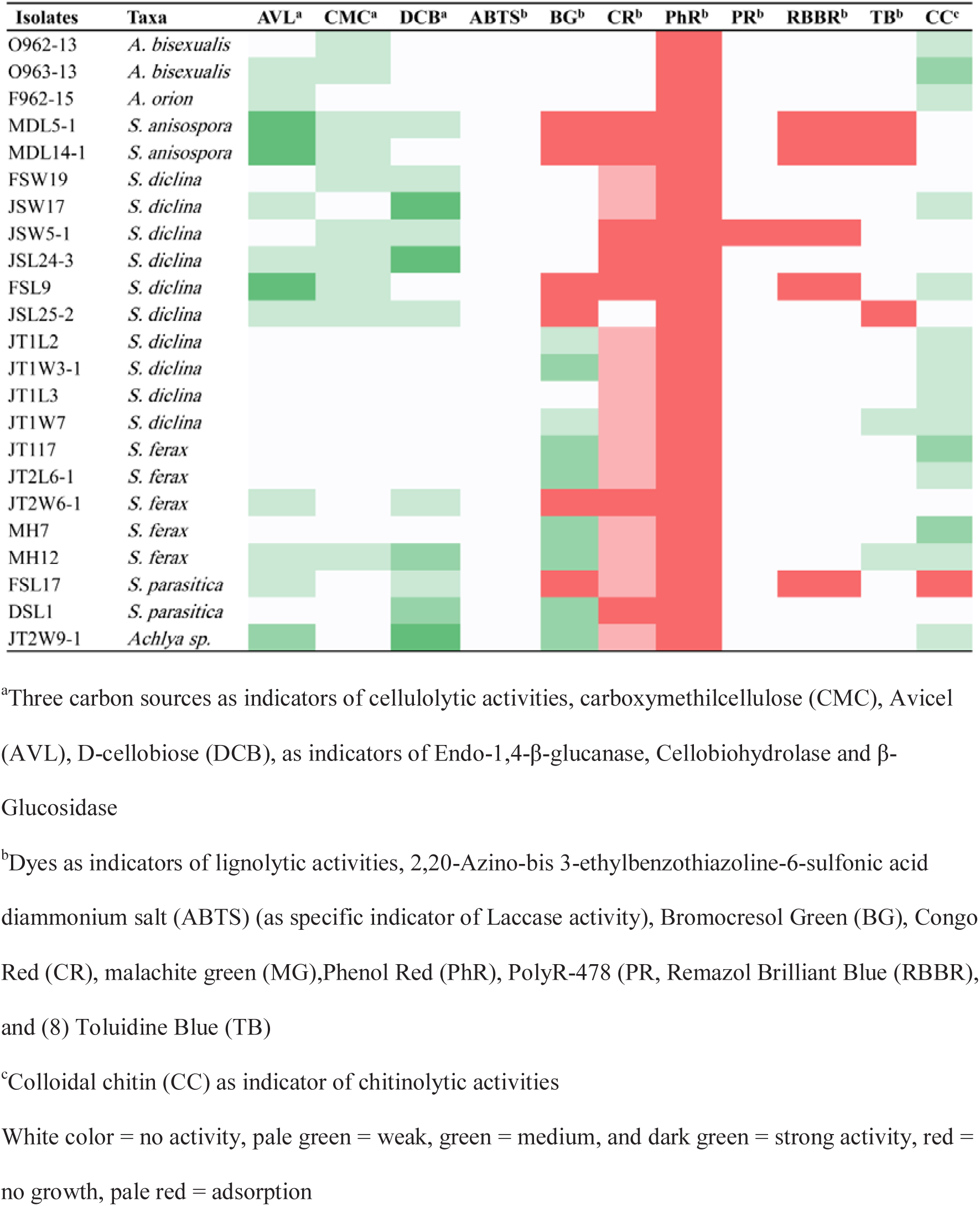
Results of experimental screening for lignolytic, cellulolytic, and chitinolytic activities of all *Saprolegniales* isolates isolated from Anzali lagoon, Rasht, Iran.

## Discussion

With this study we sought to evaluate both the spatio-temporal occurrence of *Saprolegniales* species within the brackish Anzali lagoon and to describe their morphological, phylogenetic, and physiological diversity. Previously, *Dictyuchus* Leitgeb genus (*Stramenopila, Oomycetes*) had been reported from Anzali lagoon (Masigol et al. 2018). We did not observe an influence of sampling site on the isolation of the taxa, although that isolation rate was slightly lower at the river mouth site in summer and thus may reflect some local seasonal effects. Generally, we observed a much lower abundance of *Saprolegiales* isolates in summer, when both precipitation and terrestrial input was lowest. This is in agreement with the 25 existing case studies (e.g., Czeczuga et al. 2003; Paliwal and Sati 2009) which show a similar pattern and variation in abundance. High temperature periods are the least favourable for *Saprolegniales* (Fig. 4), although this should be considered alongside other potentially covering aspects. For instance, the diversity of *Saprolegniales* isolates has been correlated with water hardness (Czeczuga et al. 2003) as well as Mg^2+^, SO_4_^2-^ and Ca^2+^concentrations (Czeczuga et al. 2002). We should also consider that the highest impact of pollution in Anzali lagoon occurs during the summer season (Fallah and Zamani-Ahmadmahmoodi 2017) associated with a high discharge of agricultural waste and increase in fish breeding activities. Khatib and Khodaparast (2010) noted that the higher water temperature and declining water volume created ideal conditions for bacteria in the summer, which might compete with oomycetes for organic matter.

**Fig. 4.**
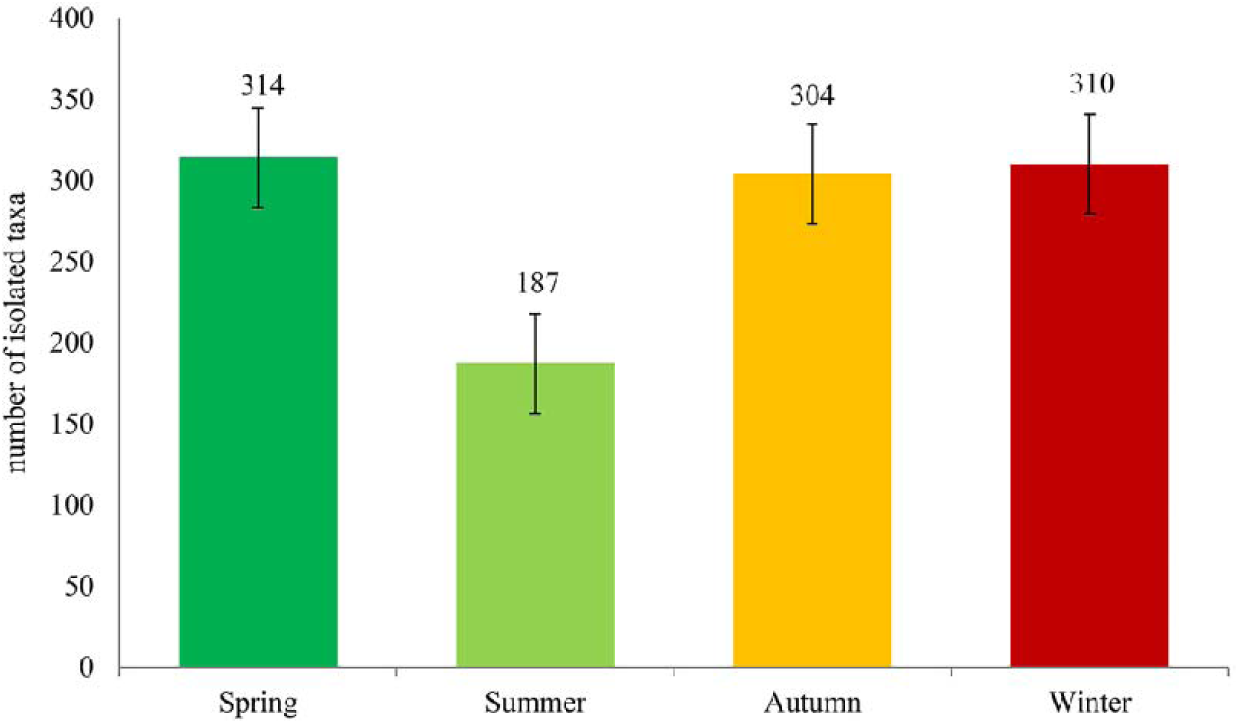
Seasonal occurrence of *Saprolegniales* isolates reported by various researchers obtained from 25 case studies (1958-2015)

Previous attempts to fully elucidate the impact of anthropogenic pollution on the microbial ecology of Anzali Lagoon have largely overlooked the role of fungi and oomycetes. Understandably, integration of mycological approaches into aquatic ecology is difficult, with vague morphological features contributing to the inability to distinguish between species *in situ*. This is particularly true for *Saprolegniales* taxa for which species identification is complex and often lacks a conclusive and consistent morphological form (Hulvey et al. 2007; Steciow et al. 2014). With this study, by assessing the number of isolated colonies we were able to better assess the occurrence and distribution of these organisms in the environment. However, this approach is still limited in that it is both time-consuming and not all *Saprolegniales* may be readily cultured.

Using a combination of morphological and molecular approaches we were able to confirm the occurrence of four *Saprolegnia* spp., of which *Saprolegnia anisospora* and *S. diclina* have never been reported in Iran. For the most part, *Saprolegnia* spp. could be differentiated using exclusively molecular approaches. *Saprolegnia anisospora*, confirmed by molecular analyses, was consistent with the morphological description albeit lacking the typical production of oospores in culture. Differentiation of *S. diclina* and *S. parasitica* and *S. aenigmatica* was only possible using molecular approaches due to a high number of shared morphological features (Johnson et al. 2002, Diéguez-Uribeondo et al. 2007, Sandoval-Sierra and Diégues-Uribeond 2015). Phylogenetic analysis generally suggested that sequences of *Achyla* spp. were similar to sequences of *Achlya bisexualis, A. flagellata* and *A. orion.* Clear approaches toward species identification are critical since many species such as *S. diclina* (Hussein et al. 2013), *S. ferax* (Cao et al. 2012) and *S. parasitica* (Griffiths et al. 2003) are considered pathogens and may devastate fish and amphibians’ populations.

As our awareness of inland waters increases over time, we are gaining an increased appreciation for their role as key players in the global carbon cycle (Tranvik et al. 2018). Inland waters, through the activity of aquatic bacteria and fungi, are involved in remineralising large proportions of terrestrial organic matter into greenhouse gases. In this context, understanding the relative contribution of bacteria and fungi, of which fungi are better equipped to break down both dissolved and particulate polymeric organic matter using adverse array of extracellular enzymes, is critical (Grinhut et al. 2011; Zahmatkesh et al. 2016; Collado 2018). However, although aquatic *Saprolegniales* are generally isolated from floated plant and animal debris, their involvement in organic matter degradation in various freshwater ecosystems has been largely ignored. In their study on seven *Saprolegniales* isolates from skin of living crayfish, Unestam (1966) showed a lack of cellulolytic activity by *Aphanomyces* spp., *Pythium* spp. and *Saprolegnia* spp. interestingly, all isolates exhibited chitinolytic activity (Unestam 1966; Nyhlen and Unestam 1975). In contrast, Thompson and Dix (1985) showed moderate to strong cellulolytic activity tested in 27 *Saprolegniales* taxa including *Achlya* spp. and *Saprolegnia* spp.. Whether the capacity of oomycetes to utilize more complex polymers, including lignin, remains unclear. Although our strains can fairly represent *Achlya* and *Saprolegnia*, isolation of less common genera will be necessary to have a better understanding of their enzymatic affinities.

In this study, capacity to degrade lignin was not evident with none of the isolates exhibiting laccase activity, and fewer than 25% of the isolates exhibiting any peroxidase activity (table 3). This agrees with a previous study where the potential for lignin degradation amongst *Dictyuchus* spp. and *Achlya* spp. isolates was essentially absent when compared to filamentous fungi isolated from the same environment (Masigol et al. 2019) and elsewhere (Abdel-Raheem and Shearer 2002; Junghanns et al. 2008; Simonis et al. 2008). The lack of any significant lignolytic activity amongst the aquatic *Saprolegniales* indicates that biopolymer degradation is specific and may be limited to just chitin and cellulose, in contrast with the broader specificity of fungi. Saprophytic *Saprolegiales* exhibiting chitinolytic and cellulolytic activity, as indicated by our study, may be more critical in the remineralisation of chitin-based particulate organic matter. This is supported by a close association of *Saprolegniales* with crustacean carapaces (Czeczuga et al. 1999; Czeczuga et al. 2002), feathers of wild and domestic bird species (Czeczuga et al. 2004), the benthic amphipod *Diporeia* spp. (Kiziewicz and Nalepa 2008) and the seeds of plants (Kiziewicz 2005) where chitin comprises a primary component of the biomass. Whilst the presence of an organism possessing chitinolytic enzymes on chitin rich substrates does not immediately prove its involvement in chitin processing, we feel it warrants additional investigation. In this study we isolated oomycetes directly from plant debris occurring in freshwater systems, despite their inability to utilise the predominantly lignin-based substrate. Steinberg et al. (2003) and Meinelt et al. (2007) argue that plant-derived humic acids, which occur at high abundance at the terrestrial-aquatic interface, inhibit the growth of some oomycetes. Therefore, their ecological role in this niche remains uncertain, although we speculate that they may form, at minimal, a commensal relationship with filamentous fungi, to utilise both the labile (chitin, cellulose) and more refractory (lignin) components on this substrate (Lennon et al. 2013; Solomon et al. 2015).

In conclusion, it is important to complement traditional morphology-based taxonomy with molecular-based taxonomy but including several markers. It will be also essential to include other techniques such as metabarcoding the have a better impression of the relative abundance of this group respect other eukaryotes. To our knowledge, most of the studies performed in various freshwaters lack precise taxonomy and hence are greatly impacting ecological interpretations. We observed clear seasonal dynamics in the occurrence of *Saprolegniales* in Anzali lagoon with a decline in summer linked to both increased water temperature and high levels of anthropogenic pollution. We confirmed that *Saprolegniales* isolates lack the broad substrate specificity of fungi, rather exhibiting specific activity towards cellulose or chitin-based substrates. Whether predominantly lignin-based plant-derived substrates are an energy source, or simply a transport vector, for aquatic oomycetes remains unclear and should be further tested. Evaluating the fate of allochthonous carbon in aquatic and global C cycles should better consider the occurrence and impact of oomycetes, particularly for substrates where chitin is particularly dominant. We should better evaluate the interactions between oomycetes and fungi and bacteria both in competition for nutrients and carbon as well as for potential commensal and synergistic impacts on carbon cycling.

